# Sound elicits stereotyped facial movements that provide a sensitive index of hearing abilities

**DOI:** 10.1101/2023.09.07.556766

**Authors:** Kameron K. Clayton, Kamryn S. Stecyk, Anna A. Guo, Anna R. Chambers, Ke Chen, Kenneth E. Hancock, Daniel B. Polley

**Author notes:** McGovern Institute for Brain Research, MIT, Cambridge, MA 02139 USA.

## Abstract

Sound elicits rapid movements of muscles in the face, ears, and eyes that protect the body from injury and trigger brain-wide internal state changes. Here, we performed quantitative facial videography from mice resting atop a piezoelectric force plate and observed that broadband sounds elicit rapid, small, and highly stereotyped movements of a facial region near the vibrissae array. Facial motion energy (FME) analysis revealed sensitivity to far lower sound levels than the acoustic startle reflex and greater reliability across trials and mice than sound-evoked pupil dilations or movement of other facial and body regions. FME tracked the low-frequency envelope of sounds and could even decode speech phonemes in varying levels of background noise with high accuracy. FME growth slopes were disproportionately steep in mice with autism risk gene mutations and noise-induced sensorineural hearing loss, providing an objective behavioral measure of sensory hyper-responsivity. Increased FME after noise-induced cochlear injury was closely associated with the emergence of excess gain in later waves of the auditory brainstem response, suggesting a midbrain contribution. Deep layer auditory cortex units were entrained to spontaneous facial movements but optogenetic suppression of cortical activity facilitated – not suppressed – sound-evoked FME, suggesting the auditory cortex is a modulator rather than a mediator of sound-evoked facial movements. These findings highlight a simple involuntary behavior that is more sensitive and integrative than other auditory reflex pathways and captures higher-order changes in sound processing from mice with inherited and acquired hearing disorders.

## INTRODUCTION

Less than one second after intense sound reaches the ear, the pupil dilates, postural muscles contract, the pinna twitches, ear drum tension increases, and inner ear molecular motors are engaged to attenuate cochlear amplification. Each of these rapid audiomotor transformations are mediated by independent neural reflex pathways that collectively protect the inner ear from acoustic injury, initiate defensive behaviors, and orient the head and body for further analysis of the sound source. In principle, assaying these involuntary behaviors with startle reflex audiometry [1], pre-pulse inhibition of startle [2,3], middle ear muscle reflex audiometry [4], olivocochlear reflex audiometry [5,6], or pupillary dilation response audiometry [7,8] provides a useful middle ground to study hearing in laboratory animals, offering higher throughput than operant behaviors while providing the integrative measure of behavioral registration in awake animals that is lacking with commonly used physiological proxies for hearing, such as the auditory brainstem response.

The primary challenge with most involuntary behavioral assays is that they are mediated by different cell types and neural pathways than the central auditory neuroaxis that provides the basis for conscious sound awareness. As reflexes that primarily serve to protect the animal and ear from injury, they are often insensitive to low and moderate sound levels that are well within the audible range and generally do not reflect the involvement of neural circuits beyond the brainstem, limiting their broader use as a behavioral measure of general hearing ability [2,9,10]. Interestingly, in analyzing high-resolution video of the face during sound presentation in mice, several recent studies observed uninstructed movements that were synchronized to the onsets of discrete sounds [11,12]. Apart from remarking on their occurrence, these studies did not delve into their acoustic feature sensitivity or characterize the neural pathways that might transform sound into facial movements.

Here, we build on these reports to show that sound-evoked movements from a region of the face just caudal to the vibrissae array are 1000 times more sensitive (30 dB) to sound than the startle reflex. We show that facial movements track the temporal envelope of sound, permitting single trial decoding of complex sounds, such as English speech tokens presented in background noise. Single unit recordings and optogenetic manipulations suggest that sound-evoked facial movements are mediated by a midbrain pathway but modulated by descending corticofugal projections. Finally, we should that facial movements capture both inherited and acquired auditory hypersensitivity in mice with mutations in autism risk genes and noise-induced cochlear sensorineural damage, respectively. In sum, our data suggest that a relatively simple approach of making video recordings of the face provides a new behavioral vantage point to study neural and behavioral processing of complex sounds.

## RESULTS

### Broadband noise bursts elicit rapid and stereotyped facial movements across a wide range of sound levels

We measured involuntary behavioral reactions to sound by placing head-fixed unanesthetized mice atop a piezoelectric force transduction plate while acquiring high-resolution video of the face. The output of the force plate provided an index of general body movements as well as large-amplitude multi-phasic events corresponding to the startle reflex (**Figure 1A**) [13]. Movement of the pinna, jaw, eyelid, nose, and pupil were quantified with DeepLabCut, a video analysis method for markerless tracking of body movements based on deep neural networks (N = 8 mice; **Figure 1B**) [14]. We also calculated facial motion energy (FME) from a region of interest caudal to the vibrissae array (**Figure 1C**), permitting simultaneous acquisition and comparison of sound-evoked movements across measurement modalities (**Figure 1D**).

**Figure 1.**
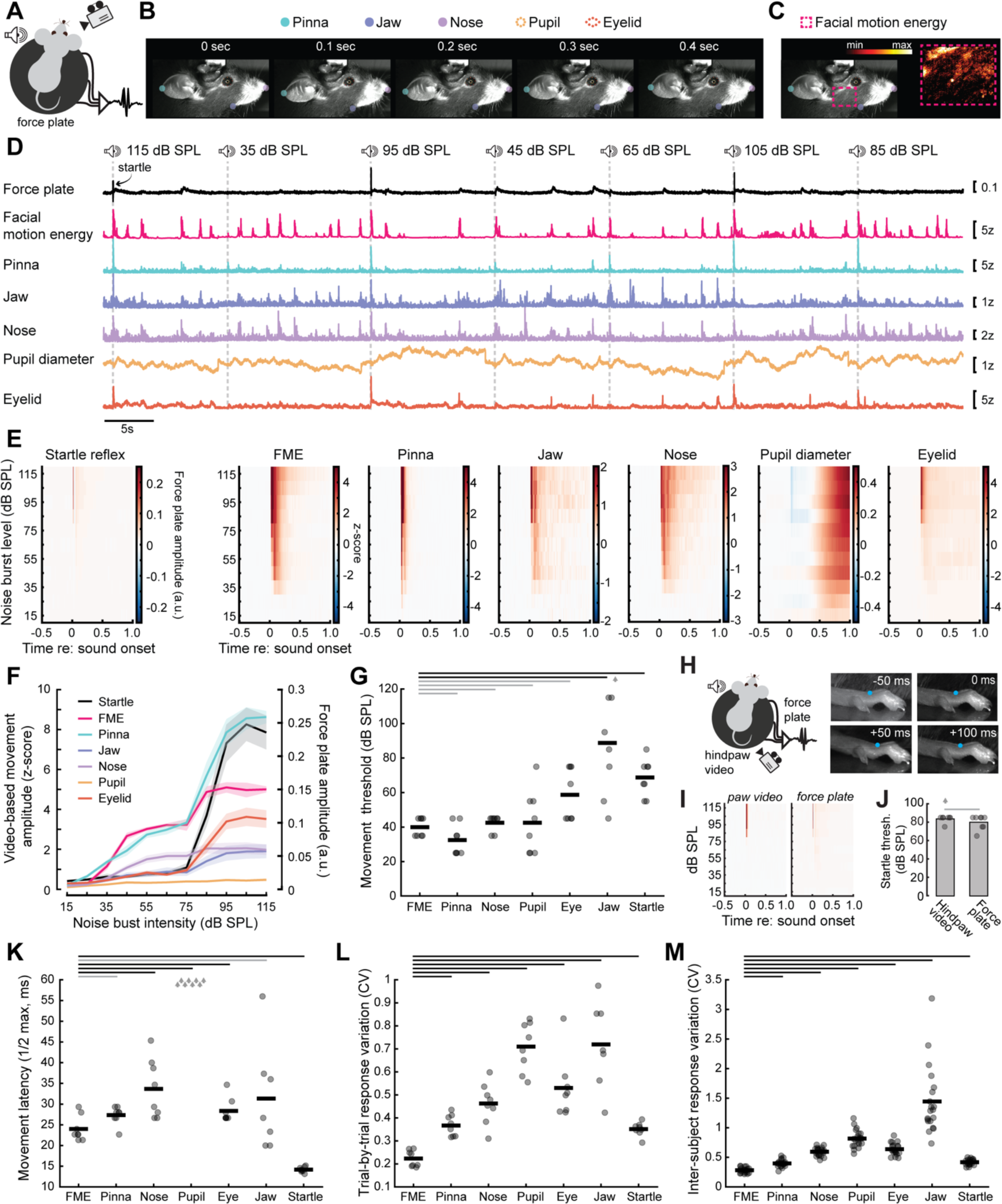
Noise bursts elicit short-latency facial movements that are 1000-times more sensitive than the acoustic startle reflex. (A) Schematic depicts mouse, camera, speaker, and summed output of three piezoelectric transducers attached to the force plate. The multi-peaked startle waveform is copied from the first 115dB SPL trial shown in *D*. (B) Video frames from a single trial depict changes in pinna, jaw, nose, pupil, and eyelid positions determined by DeepLabCut. (C) Motion energy calculated from a region of interest positioned caudal to the vibrissa array. (D) Movement amplitudes for 7 consecutive trials in a representative mouse. Y-axis scales presented at right. (E) Mean peristimulus force plate and facial movement responses from 8 mice. (F) Mean ± SEM movement amplitudes. Vertical axes on the left and right refers to the videographic movement and startle reflex, respectively. (G) Minimum sound intensity that elicited movement presented for each mouse (circle) and sample mean (horizontal bar). Threshold varied significantly across movement types (one-way repeated measures ANOVA, F = 13.45, p = 2 × 10^−8^) with post-hoc comparisons finding significant differences between FME and jaw (p = 0.01) and startle reflex (p = 0.0002). For all figures, black and gray horizontal bars denote significant (p < 0.05) and non-significant differences, respectively, with FME after Holm-Bonferroni corrections for multiple comparisons. (H) *Left:* As per *A*, except that high-resolution video recordings (150 frames/s) are made of the hindpaw. *Right:* Blue circle indicates DeepLabCut tracking of a point on the hindpaw in four frames relative to the onset of a noise burst. (I) Mean peristimulus force plate and hindpaw video responses from 8 mice. Force plate and hindpaw pseudocolor scales match the startle reflex and nose videography plots presented above in *E*. (J) Startle thresholds for each mouse (circle) and sample mean. Arrow indicates one outlying value outside of the plotted range. Gray bar indicates non-significant difference in startle reflex threshold measured via the piezoelectric force plate or hindpaw videography (paired t-test, p = 0.17). (K) Sound-evoked movement latencies presented for each mouse (circle) and sample mean (horizontal bar). Response latency varied significantly across movement types (one-way repeated measures ANOVA, F = 550.67, p = 8 × 10^−34^) with post-hoc comparisons finding significant differences between FME and nose (p = 0.01), pupil dilation (p = 2 × 10^−7^), eyelid (p = 0.01), and startle reflex (p = 0.0003). Gray arrows for pupil denote that values were outside of the y-axis range (mean latency = 627 ms). (L) Trial-to-trial variability measured with the coefficient of variation presented for each mouse (circle) and sample mean (horizontal bar). Trial-to-trial variability was significantly different between movement types (one-way repeated measures ANOVA, F = 23.18, p = 6 × 10^−11^) with post-hoc comparisons finding that FME was significantly less variable than all other movement types (p < 0.0004 for all comparisons). (M) Inter-subject variability measured with the coefficient of variation across subjects for each trial (circle) and sample mean (horizontal bar). Inter-subject variability was significantly different between movement types (one-way repeated measures ANOVA, F = 59.12, p = 1 × 10^−34^) with post-hoc comparisons finding that FME was significantly less variable than all other movement types (p < 0.005 for all comparisons).

We observed that broadband noise bursts elicited short-latency facial twitches and longer latency pupil dilations across a far wider range of sound levels than the acoustic startle reflex (**Figure 1E**). To determine whether apparent differences between facial movements and startle could be attributed do inherent sensitivity differences between measurement modalities, we also performed high-speed videography of the hindpaw in a subset of mice but noted the close correspondence between force plate and hindpaw movement amplitudes (N = 8; **Figure 1F**). To quantify differences in the amplitude of these movements across the full range of sound levels, z-scored amplitudes of videographic measures were compared with changes in force plate amplitude. Acoustic startle reflex amplitudes increased rapidly above 75 dB SPL and then saturated at 95 dB SPL (**Figure 1G**). Sound-evoked movement of the eyelid and jaw showed the same high-threshold saturating response as the acoustic startle response, while the pupil was less responsive overall than other facial markers. By contrast, FME, as well as movement of the pinna and nose, grew monotonically across a far wider range of sound levels, suggesting that they may arise from a separate pathway for neural sound processing than the acoustic startle reflex.

Using FME as a point of reference to other movement markers, we found that FME thresholds were approximately 40 dB SPL and not significantly different than movement thresholds for the pinna, nose, pupil, or eyelid but were significantly more sensitive than the startle reflex or jaw movements (**Figure 1G**, statistical reporting provided in the figure legends throughout). To control for the possibility that startle thresholds measured with the piezoelectric force plate were inherently less sensitive than videography, we performed an additional experiment featuring combined measurements of the force plate with high-resolution videography of the hindpaw (**Figure 1H)**. Both measurements showed similarly brief, high-threshold responses to the same broadband noise stimulus described above (**Figure 1I-J**), underscoring that the startle reflex – regardless of measurement modality – is distinct from sound-evoked facial movements.

Facial movements were surprisingly fast, occurring less than 50 ms following sound onset (**Figure 1K**). The startle reflex was fastest of all (14.2 ± 0.2 ms), while sound-evoked pupil dilations were over an order of magnitude slower than other movements, with an average onset latency of 627.3 ± 22.8 ms. Although FME, pinna, and nose movements showed comparable thresholds and latencies, FME proved the most robust measure of sound-evoked facial movements, as evidenced by significantly lower trial-by-trial variability (**Figure 1L**) and inter-subject variability (**Figure 1M**) than all other movement markers. For this reason, we relied on FME for all subsequent analyses.

### Facial movements track the temporal envelope of broadband sounds and provide an index of speech processing in background noise

Most commonly used reflexive and physiological markers of sound processing rely on discrete bursts of sounds to elicit responses of varying magnitude. However, encoding natural sounds is critically depending on tracking fluctuations in the sound pressure envelope over time [15]. Hence, an involuntary behavioral readout of sound processing that indexed synchronization to the sound pressure envelope could provide insights into neural encoding of more complex sounds that are not possible with other reflexive or voluntary behaviors.

To address the possibility that sound-evoked facial movements could provide this insight, we first introduced silent gaps of varying duration in continuous broadband noise. We found that facial movements were elicited at the offset of silent gaps (**Figure 2A**), where the magnitude of FME grew monotonically with gap duration (**Figure 2B**), with FME gap response thresholds of approximately 55 ms across eight mice (**Figure 2C**). Next, we noted that facial movements were entrained to the repetition rate of broadband sounds, such as FM sweeps (**Figure 2D**). We quantified facial synchronization to the sound pressure envelope by performing a Fourier analysis on FME amplitude over time and calculating the power at the stimulus repetition rate relative to the noise floor (**Figure 2E**). We found that FME became less synchronized as the presentation rate of FM sweeps increased but tracked the sound envelope for modulation rates up to 3 Hz (**Figure 2F**).

**Figure 2.**
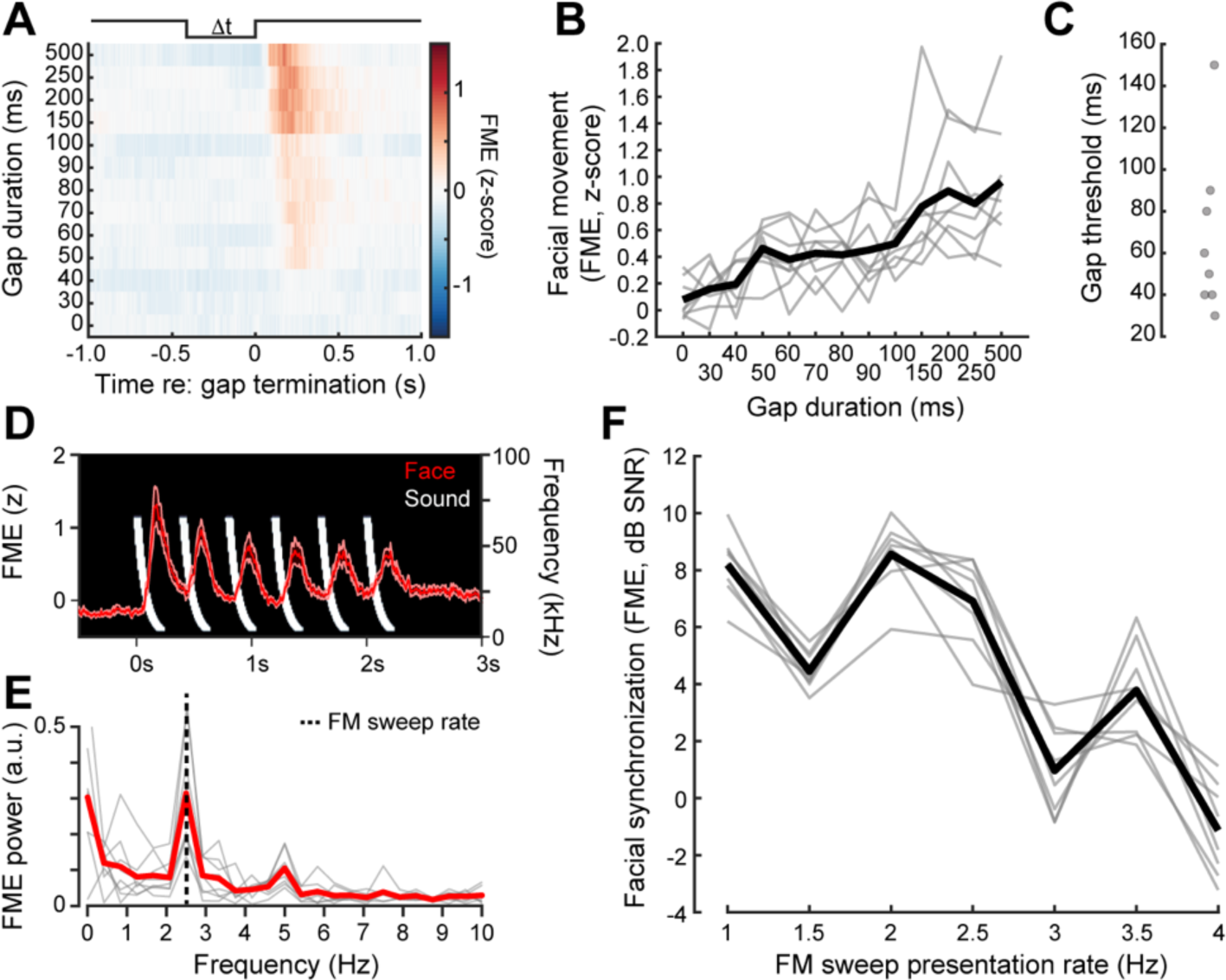
Facial movements synchronize to slow changes in the sound pressure envelope. (A) Silent gaps of varying durations were introduced in a constant background of 50 dB SPL white noise. Mean FME before, during, and after the silent gap (N = 8). (B) FME significantly increased with gap duration (one-way repeated measures ANOVA, F = 9.43, p = 3 × 10^−11^). Individual mice and sample mean (N=8) are plotted as thin gray lines and thick black line, respectively. (C) Gap detection thresholds for 8 mice. (D) Spectrogram depicts downward frequency modulated sweeps presented at 2.5 Hz with a 50% duty cycle (white) at 70 dB SPL. Mean ± SEM FME amplitude for an example mouse (red) shows a facial twitch elicited by each of the six consecutive FM sweeps. (E) Fourier analysis of FME responses from 8 mice to the FM sweep sequence presented at 2.5Hz reveals a peak at the presentation rate (dashed vertical line). Individual mice and sample mean are plotted as thin gray lines and thick red line, respectively. (F) Facial synchronization was calculated as the power at the stimulus presentation rate relative to the noise floor. Synchronization significantly decreases across higher FM sweep presentation rates (one-way repeated measures ANOVA, F = 52.73, p = 6 × 10^−18^).

In the hearing sciences, measures of response thresholds in silence are often poorly predictive of complex sound processing in background noise [16,17]. Understanding the broader impact of hearing loss or hearing restoration interventions on behavioral registration of communication sounds remains a high priority for animal models of human hearing disorders. Therefore, as a next step, we presented sequences of two English phonemes (*Gee* and *Ha,* 70 dB SPL at 1 Hz, **Figure 3A-B**) and quantified facial movement synchronization to these repeating speech tokens in increasing levels of background noise (**Figure 3C**). We found that the synchronization to the speech presentation rate decreased with background noise level but remained fairly robust up to 50 dB SPL background noise (**Figure 3D**).

**Figure 3.**
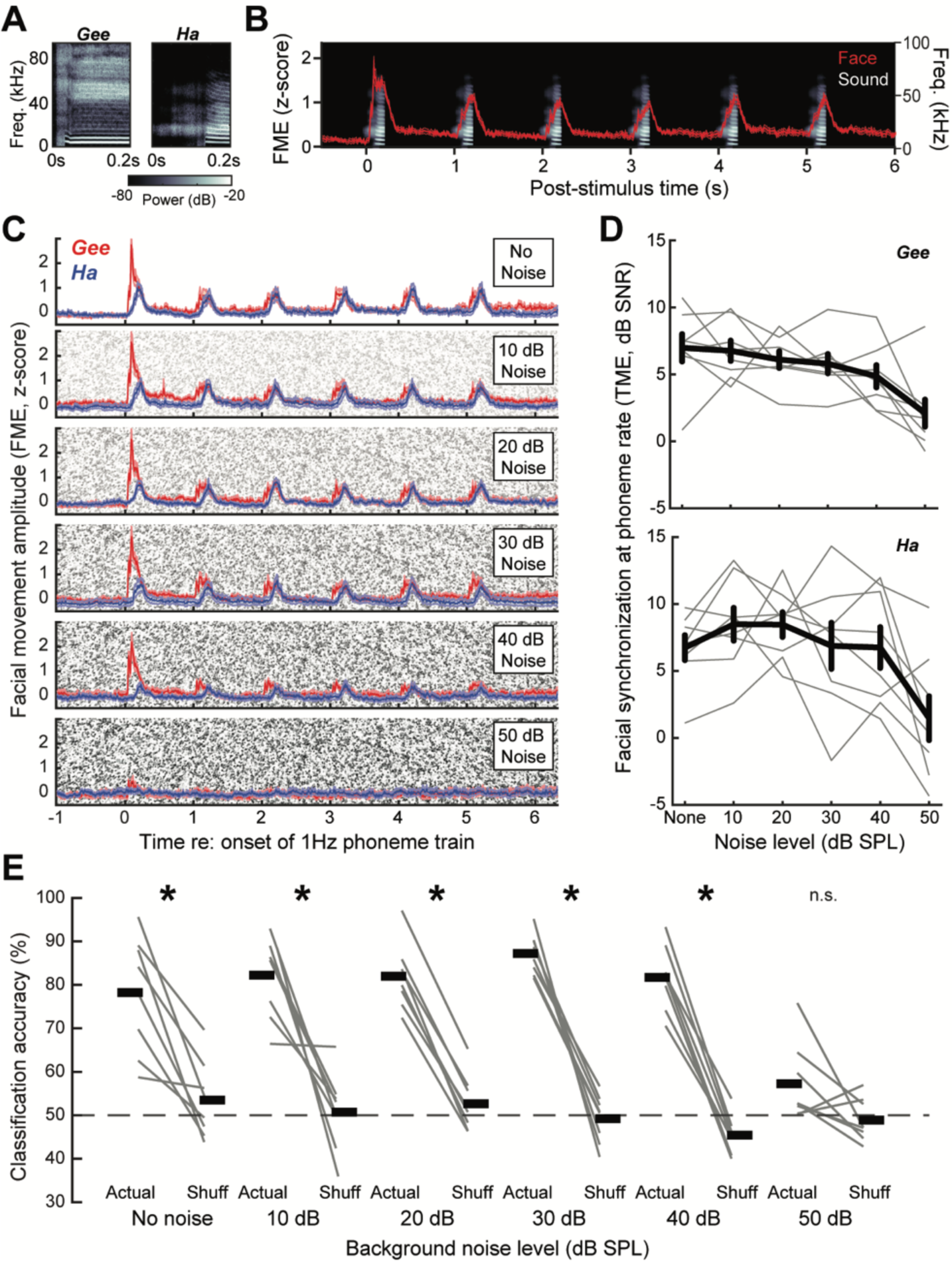
Decoding phonemes in background noise via facial movements. (A) Spectrograms of two English speech tokens digitally resynthesized to span the mouse hearing range without distorting the spectrotemporal envelope of the source signal. (B) Spectrogram plots six presentations of the phoneme, *Gee,* presented at 1Hz (grayscale, right vertical axis). Mean ± SEM FME from a representative mouse elicited by each speech token (red, left vertical axis). (C) Mean ± SEM FME for *Gee* and *Ha* presented without background noise and five levels of increasingly intense background noise (N = 8). (D) FME synchronization to the speech token presentation rate decreased significantly, but equivalently, for both phonemes across increasingly levels of background noise (2-way repeated measures ANOVA, main effect for noise level [F = 12.36, p = 2 × 10^−8^], main effect for phoneme [F = 0.78, p = 0.39]). (E) Single trial speech token classification accuracy with actual and shuffled (shuff) assignment of stimulus identity. Chance classification = 50%. Classification accuracy was significantly greater for actual than shuffled stimulus label assignments for all background noise levels (paired t-tests, p < 0.002 for all), except at the highest noise level (50 dB SPL, p = 0.07).

Measuring whether a speech stimulus can be detected in noise is less relevant than whether it can be discriminated in noise. The two tokens presented clearly differed in spectrum and voice onset timing (Figure 3A), such that the phase of facial synchronization to *Gee* and *Ha* was slightly offset (see timing of blue versus red movements, Figure 3C). To determine whether these timing cues in sound-evoked facial movements provided a basis for classifying the speech token presented in varying levels of background noise, we performed a principal components analysis to decompose the movement vectors into a lower dimensional space and then trained a support vector machine to classify held-out trials as either *Gee* or *Ha*. We found approximately 80% speech token classification accuracy, which was significantly higher than chance for all noise levels up to 50 dB SPL (**Figure 3E**). These findings demonstrate that a video-based analysis of involuntary facial twitches in mice can provide insight the neural processing of complex communication sounds that is not possible with other commonly used measures of hearing in laboratory animals.

### Facial movements are insensitive to pure tones

Facial videography offers several advantages over traditional indices of hearing function like the ABR, in that it can be measured in unanesthetized animals and provides a direct behavioral readout for the encoding of spectrotemporally complex sounds instead of relying only on short tone bursts or clicks. The principal use of ABR measurements is to provide a non-invasive physiological assay of cochlear function. For example, several hours of exposure to a 103 dB SPL 16-32 kHz noise band damages outer hair cells and cochlear afferent synapses in the high-frequency base (**Figure 4A**). The sensorineural damage can be directly visualized with post-mortem cochlear histology [18] but can be indirectly indexed in vivo by a permanent increase in ABR thresholds evoked by tone bursts 16 kHz and higher (**Figure 4B**). To determine whether sound-evoked facial movements can also provide an index of cochlear sensorineural hearing loss (SNHL), we measured FME evoked by pure tones (**Figure 4C**) but discovered that facial movements were significantly less responsive to tones than broadband noise, with average response thresholds as high as 90 dB SPL (**Figure 4D**).

**Figure 4.**
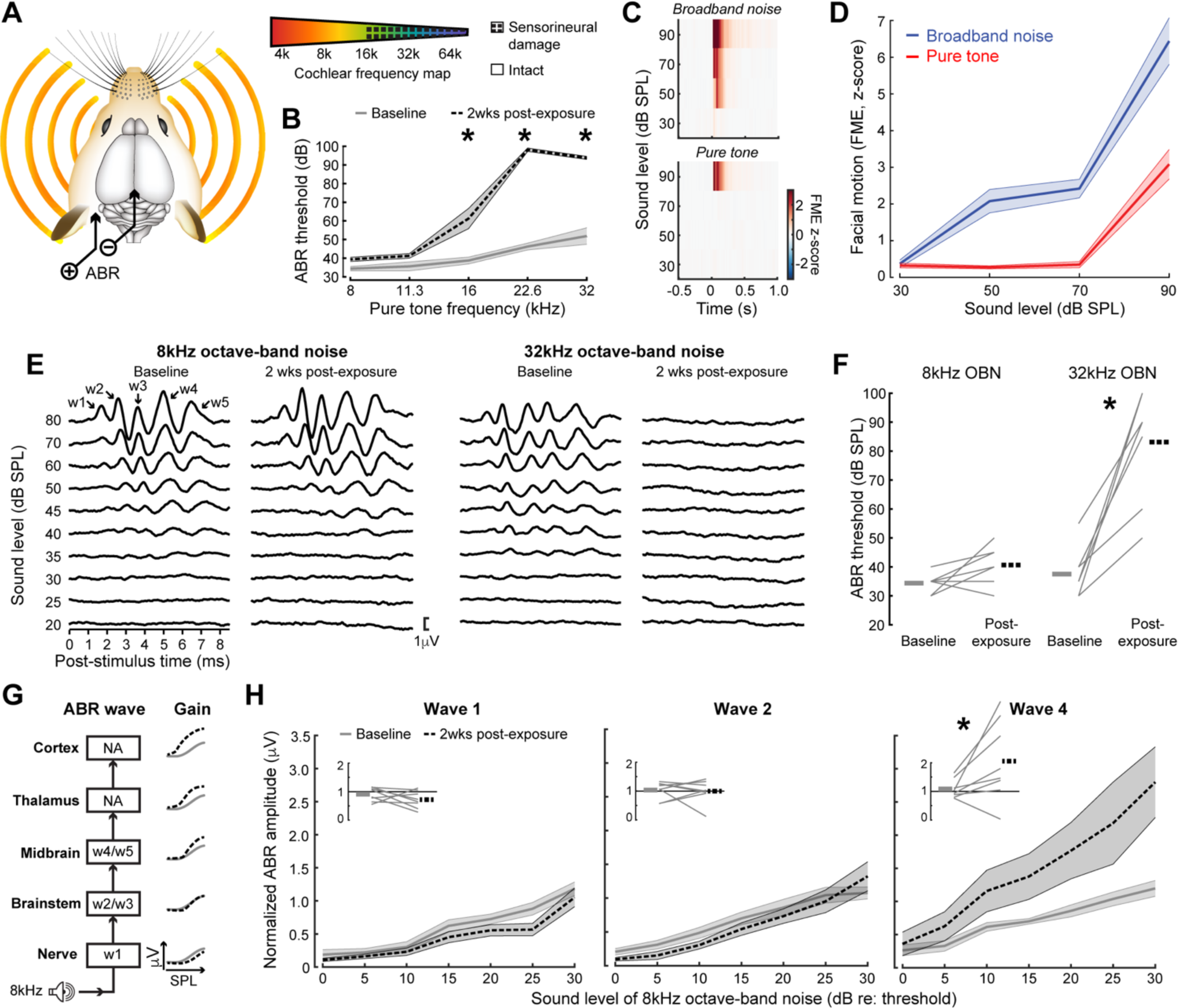
Noise-induced SNHL causes a combination of high-frequency threshold shift and excess low-frequency gain in the ABR. (A) Two hours of exposure to 16-32 kHz octave-band noise at 103 dB SPL causes sensorineural damage in the high-frequency base of the cochlea, as determined with pinna-vertex ABR measurements. (B) Tone-evoked ABR measurements confirm threshold elevation for test frequencies above 11.3 kHz measured before vs two weeks after noise exposure (2-way repeated measures ANOVA, N=9, main effect for group, F = 146.95, p = 9 × 10^−9^; group × frequency interaction, F = 39.73, p = 9 × 10^−16^]). Asterisks denote post-hoc pairwise comparisons (p < 0.05 for all). (C) Mean peristimulus FME responses from 8 mice for pure tone and broadband noise stimuli. (D) Mean ± SEM FME amplitudes over sound levels. Broadband noise evokes significantly larger facial movements than pure tones [2-way repeated measures ANOVA, N = 8, main effect for stimulus type, F = 54.95, p = 4 × 10^−6^; group × frequency interaction, F = 10.87, p = 3 × 10^−5^]. (E) Octave-band noise centered at 8 and 32kHz elicits a robust multi-peaked ABR before acoustic trauma. Two weeks after noise-induced high-frequency SNHL, the ABR response to 32kHz noise band is virtually absent at sound levels up to 80 dB SPL, whereas responses to the 8 kHz noise band appear unaffected or slightly larger than baseline measurements. Arrows indicate the appearance of ABR waves (w) 1-5. (F) Octave-band noise ABR thresholds after noise exposure (two-way repeated measures ANOVA, N = 8; main effect for frequency [F = 38.26, p = 2 × 10^−5^]; main effect for timepoint [F = 46.59, p = 8 × 10^−6^], frequency × timepoint [F = 38.259, p = 1 × 10^−4^]). (G) Schematic illustrating the neural generators of ABR waves 1-5 and the expected transition from slight attenuation in the 8 kHz OBN level × amplitude input-output function for early waves to excess central gain measured in later waves. NA indicates that the neural generators of the ABR do not include central auditory structures above the midbrain. (H) Mean ± SEM 8 kHz OBN-evoked normalized wave amplitude growth functions plotted relative to threshold. *Inset:* 8 kHz OBN-evoked normalized wave amplitudes were averaged within a 30-45 dB range above threshold. Solid black line represents no change, gray lines represent individual subjects, thick gray and dashed black lines represent baseline and 2 weeks post-exposure, respectively. Asterisks denote p < 0.05.

### ABR and facial movements provide congruent measures of hearing loss and excess central gain after acoustic trauma

To address whether the stimuli used for ABR and facial videography measures could be adapted to support direct comparison, we tested a stimulus with a spectral bandwidth in between a pure tone and broadband noise. We measured the ABR with octave-band noise centered on 8 or 32 kHz and observed that both stimuli elicited low-threshold responses with the expected multi-peaked waveform in a baseline measurement session prior to noise exposure (**Figure 4E**). Two weeks after high-frequency noise exposure, we observed that responses remained robust at 8 kHz but were strongly attenuated at 32 kHz, with thresholds elevated by approximately 40 dB, as observed with pure tone stimuli (**Figure 4F**).

Exposure to intense noise has two types of effects on sound-evoked neural activity: First, neural responses to high stimulus frequencies within the range of cochlear damage are reduced due to degeneration of cochlear sensory cells and primary afferent nerve endings [19]; Second, single unit responses to lower stimulus frequencies bordering the cochlear lesion are unaffected at early stages of auditory processing but are enhanced at higher stages of the central auditory pathway due to the increased expression of compensatory plasticity processes that are collectively described as excess central gain (**Figure 4G**) [18,20–24]. To determine whether excess central gain becomes more prevalent at successive stages of neural processing with gross neuroelectric recordings, we analyzed the amplitude of each ABR wave before and after noise exposure. Each wave of the ABR is generated by the initial volley of synchronized action potentials at successive stages of central auditory processing [25,26]. Thus, wave 1 is generated by spiral ganglion neurons, waves 2 and 3 are generated by the cochlear nucleus and superior olivary complex, and waves 4 and 5 are generated within the auditory midbrain [27–30]. The 8 kHz octave-band noise reliably elicited waves 1, 2, and 4 across our subjects, affording us the opportunity to quantify the amplitude growth for each wave across a fixed range of sound levels before and after noise exposure. As predicted, we observed that normalized input-output functions for the low-frequency stimulus were slightly attenuated for wave 1, unchanged from baseline for wave 2, but were enhanced above baseline for wave 4, consistent with the emergence of central compensatory plasticity mechanisms at higher stages of central auditory processing (**Figure 4H**).

We found that 8 and 32 kHz octave-band noise elicited strong facial movements (**Figure 5A**) that increased over a wide range of sound levels (**Figure 5B**) and remained stable over a 2-week measurement period following an innocuous sham noise exposure (**Figure 5C**). As expected from ABR threshold changes (Figure 4B), response thresholds for 8 kHz octave-band noise did not change after intense noise exposure that caused high-frequency SNHL, while 32 kHz responses were elevated by 40-50 dB (**Figure 5D**). Based on reports that central gain emerges earlier and is stronger at later stages of the central pathway following cochlear afferent lesions [20,23,31], we hypothesized that if sound-evoked facial movements were mediated by a brainstem pathway we would – like the early waves of the ABR – observe an invariant 8 kHz-evoked response amplitude over the measurement period and stably depressed 32 kHz-evoked responses (**Figure 5E**). Conversely, if sound-evoked reflexes were mediated by auditory midbrain or forebrain nuclei, we would expect that 8 kHz-evoked facial movements would grow to exceed baseline levels within days (forebrain) or weeks (midbrain) following high-frequency SNHL, while 32 kHz-evoked movements would be initially reduced before staging a partial recovery to baseline levels.

**Figure 5.**
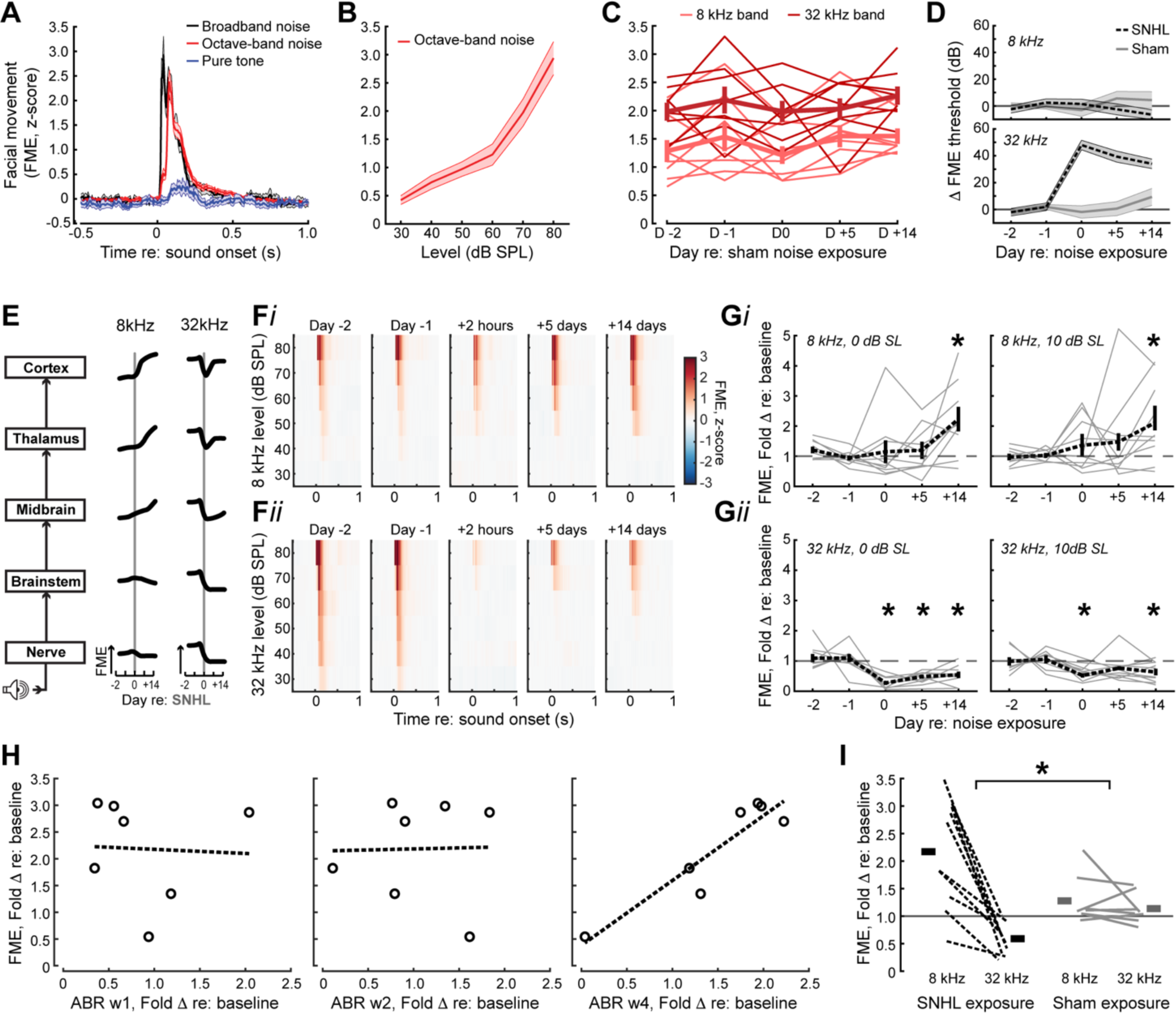
Changes in sound-evoked facial movement after noise-induced SNHL parallel modifications in late ABR waves. (A) Mean ± SEM facial movements evoked by broadband noise, octave-band noise, and pure tones at 70 dB SPL. Octave-band noise and pure tone responses are averaged at 8 and 32 kHz. (B) Octave-band noise responses are elicited at low sound levels and grow monotonically with sound level (One-way repeated measures ANOVA [F=189.9, p = 6 × 10^−66^, N = 20]). (C) Sound-evoked facial movements elicited by octave-band noise centered at 8 and 32 kHz are stable over a 17 day (D) measurement period spanning sham noise exposure (2-way repeated measures ANOVA, N = 8, main effect for Day, [F = 1.2, p = 0.33]; main effect for Frequency, [F = 40.1, p = 4 × 10^−4^]). (D) *Top:* Thresholds for sound-evoked facial movements with the 8kHz octave-band noise are unchanged over time after noise exposure and do not differ between SNHL and sham groups (2-way repeated measures ANOVA, N = 18, main effect for Day [F = 0.14, p = 0.93]; main effect for Group [F = 0.44, p = 0.52]). *Bottom:* Thresholds for sound-evoked facial movements with the 32 kHz octave-band noise are elevated after SNHL but not sham noise exposure (2-way repeated measures ANOVA, N = 18, main effect for Day, [F = 32.87, p = 1 × 10^−11^]; main effect for Group, [F = 39, p = 1 × 10^−5^]). (E) Schematic illustrating hypothetical changes in sound-evoked facial movement amplitudes over the same 17-day period before and after a SNHL-inducing noise exposure (vertical gray line). The cartoon model assumes central gain is progressively enhanced at successive stages of the central pathway, promoting faster and more complete recovery of the high-frequency response in the damaged region of the cochlea and hyper-responsiveness to the low-frequency noise band. (F) Mean peristimulus FME responses from 8 mice over a 17-day period spanning noise-induced hearing loss for an 8 kHz octave-band noise (bottom) and 32 kHz octave-band noise (bottom). (G) Actual data after SNHL are compared against the cartoon model shown in *E* by plotting the fold change in facial movement amplitudes relative to baseline (mean of D-1 and D-2) for sound levels at threshold or 10 dB above threshold (i.e., 0dB and 10dB sensation level, SL). Data from individual mice (N=10) and group mean are shown as thin gray and thick dashed lines, respectively. Asterisks denote that the change in sound-evoked facial movements are either significantly elevated relative to baseline (top) or significantly suppressed relative to baseline (bottom), as assessed with one-sample t-tests relative to a population mean of 1.0 (p < 0.02 for all significant time points). (H) Changes in ABR amplitude and sound-evoked facial movements elicited by the 8 kHz noise band after noise-induced high-frequency SNHL are calculated for a subset of mice with data from both measurement types (N = 7). Dashed line presents the linear fit of the data. Facial movement changes reflect the average of 0 and 10dB SL sound levels. Increased 8kHz-evoked facial movements are not correlated with changes in ABR wave 1 (Pearson r = −0.05, p = 0.92) or ABR wave 2 (r = 0.02, p = 0.96), but is significantly correlated with changes in ABR wave 4 (r = 0.92, p = 0.003). (I) The pattern of increased responses at 8 kHz and decreased responses at 32 kHz observed two weeks after SNHL is not observed following sham exposure (Two-way mixed model ANOVA, main effect for Group [F = 0.65, p = 0.42]; main effect for Frequency [F = 22.45, p = 3 × 10^−4^]; Group × Frequency interaction [F = 15.65, p = 0.001]). Thin and thick lines represent data from individual mice and group means from the SNHL group (N = 10) and sham exposure group (N = 8), respectively.

Sound-evoked facial movements exhibited a pattern of loss and recovery after SNHL most consistent with a neural pathway including midbrain or forebrain nuclei (**Figure 5F**). We found that facial movements evoked by the spared 8 kHz noise band at near-threshold intensities were initially stable after noise exposure but then became significantly greater than baseline levels 2 weeks following noise exposure (**Figure 5G**, *top*). Similarly, facial movements evoked by the 32 kHz noise band were nearly eliminated hours after noise exposure but were partially recovered 2 weeks after noise exposure (**Figure 5G**, *bottom*). Comparing changes in the 8 kHz ABR and FME amplitudes in the same mice 2 weeks after noise-induced SNHL, we observed that mice with the strongest enhancement of FME amplitude also had the strongest increase in wave 4 amplitude, but observed no correlation between FME and changes in the amplitude of waves 1 or 2 (**Figure 5H**). Finally, the combination of enhanced facial movements at 8 kHz and suppressed responses at 32 kHz observed after SNHL was not a consequence of repeated testing over the 2-week period, as facial movement amplitudes at both test frequencies remained relatively stable in sham-exposed mice (**Figure 5I**).

### Suppressing auditory cortex neural activity enhances sound-evoked facial movements

To study the correspondence between facial movements and neural activity dynamics in brain regions downstream of the auditory midbrain, we performed extracellular single unit recordings from the primary auditory cortex (A1) with 64-channel laminar probes (**Figure 6A**). Isolated regular spiking (RS) single units increased their activity shortly after the presentation of broadband sounds, though it is difficult to isolate the relative contribution of sound-evoked sensory inputs from motor-preparatory inputs in instances where they overlap in time. For this reason, we identified spontaneous movements that occurred in the absence of sound (**Figure 6B**). We identified the layer (L) 4/5 boundary from the appearance of a short-latency sink in the translaminar current source density (**Figure 6A**, white arrow) and assigned each RS unit to L2/3, L4, L5, or L6. In L2/3 and L4, only 11% of RS units exhibited significant spike rate modulation during spontaneous facial movements (all positively modulated), whereas 26% of units in L5 and L6 were significantly modulated (17% positively modulated), (**Figure 6C-D**). A more granular analysis of spike rate modulation by depth confirmed prior reports that deep layer auditory cortex (ACtx) neurons are more strongly recruited by movement (**Figure 6E**) [32–35].

**Figure 6.**
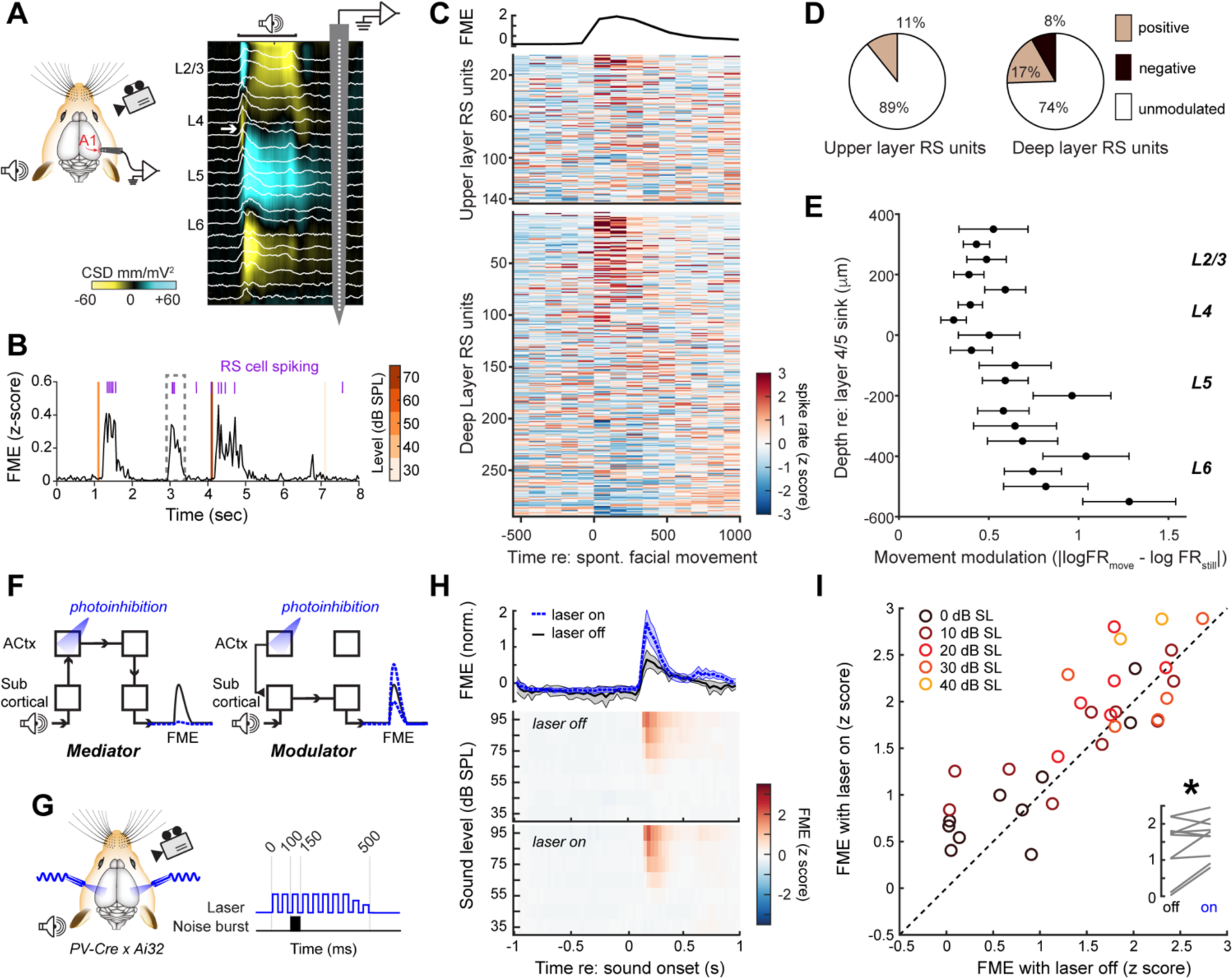
Suppressing auditory cortex activity facilitates sound-evoked facial movements. (A) Extracellular recordings were made from all layers of the primary auditory cortex (A1) with a 64-channel linear probe during contralateral sound presentation and facial videography. Electrophysiological responses are filtered offline to separate spiking activity (white trace) and the current source density (CSD). White arrow in CSD trace identifies the early current sink in layer (L) 4 elicited by a 70 dB SPL 50 ms white noise burst that is used to assign units to layers. (B) FME is increased following sound presentation (orange line) but facial twitches also occur spontaneously (dashed box). Action potentials (purple) from a single regular spiking (RS) unit are evoked by the combination of sound and movement but also during spontaneous facial movements. (C) Neurograms present the z-scored firing rates before and after bouts of spontaneous facial movements from 438 single RS units grouped into superficial (L2/3 and L4) and deep (L5 and L6) layers of the cortical column. Units are sorted by their mean activity. The line plot presents the mean FME over the same period. (D) Pie charts represent whether and how single unit firing rates were modulated by spontaneous facial movements. (E) Mean ± SEM absolute value of firing rate changes during spontaneous facial movements along the cortical column. Spike rate modulation is significantly elevated with increasing depth in the cortical column (one-way ANOVA, n = 438, main effect of depth [F = 2.09, p = 0.006]). (F) *Left:* Schematic illustrates that optogenetic suppression of auditory cortex (ACtx) spiking could eliminate sound-evoked facial movements if it were an obligatory sensorimotor relay. *Right*: Alternatively, optogenetic suppression of ACtx spiking could amplify or attenuate sound-evoked facial movements if it modulated a subcortical sensorimotor relay. (G) Illustration of experimental paradigm to test the two hypothetical scenarios described above in a transgenic mouse that expresses channelrhodopsin in parvalbumin-expressing GABAergic interneurons (PVIs). Trials of pulsed bilateral activation of ACtx PVIs throughout the peristimulus period were interleaved with sound-only trials. (H) Sound-evoked FME on interleaved trials of ACtx suppression (laser on) or no suppression (laser off). *Top:* Mean ± SEM FME. *Bottom:* Mean FME over a range of broadband noise levels (N = 10). (I) Scatterplot presents mean sound-evoked FME during ACtx suppression versus no suppression. Each symbol represents the trial-averaged mean from a single animal color coded according to the level of the sound relative to threshold. Data points above the line of unity (dashed diagonal) are enhanced during ACtx suppression. *Inset*: mean FME within 10 dB SPL above threshold is plotted for each mouse during laser off and on trials. Asterisk denotes significant difference (paired t-test, p = 0.005).

Deep layer ACtx neurons respond both to sound and movement, suggesting that they could be an obligatory relay for converting the sensory representation into a motor signal. On the other hand, sound-evoked first spike latencies are on the order of 15-25 ms in deep layer ACtx neurons [36], which is approximately coincident with the sound-evoked FME response (24 ± 1.1ms; Figure 1H). By contrast, sound-evoked first spike latencies in the inferior colliculus, an auditory midbrain structure, are on the order of 6-10 ms, thus occurring well ahead of sound-evoked movements [37]. Inferior colliculus neurons also generate wave 4 of the ABR, which showed a close correspondence with changes in sound-evoked facial movements after noise exposure (**Figure 5H**) and also receive massive corticofugal feedback from the ACtx. Collectively, these pieces of evidence suggest an alternative model where ACtx is not a mediator of sound-evoked facial movements but instead modulates the sensorimotor transformation via its descending projections to subcortical nuclei.

We reasoned that it would be possible to test these competing models by suppressing ACtx RS spiking and comparing sound-evoked facial movements with and without ACtx photoinhibition. If ACtx mediated the response, sound-evoked facial movements would be eliminated during cortical photoinhibition (**Figure 6F**, *left*). If ACtx played a modulatory role, sound-evoked facial movements would be attenuated or enhanced during cortical photoinhibition (**Figure 6F**, *right*). We suppressed ACtx spiking via bilateral optogenetic activation of parvalbumin-expressing GABAergic interneurons before, during, and after the presentation of a broadband noise stimulus and interleaving laser on trials with laser off trials (**Figure 6G**). Our findings clearly supported a modulatory model by demonstrating significant facilitation of sound-evoked facial movements on trials with optogenetic activation of GABAergic interneurons (**Figure 6H-I**).

### Sound-evoked facial movements capture an auditory sensory hyper-responsivity phenotype in mice with an autism risk gene mutation

Sensory overload is a cardinal feature of autism spectrum disorder, hyperacusis, schizophrenia, traumatic brain injury, post-traumatic stress disorder, and other neurodevelopmental disorders. Overload is particularly acute in the auditory modality, where subjects often report that moderate intensity sounds are uncomfortably loud, unpleasant, or even painful [38]. The sensory overload phenotype can be challenging to model in laboratory animals with mutations in risk genes for these conditions because the broader cognitive and motor impairment can interfere with learning the procedural demands of an operant behavioral task. Here, we focused on mice with deletion of PTCHD1, an × chromosome gene found in families with autism spectrum disorder and intellectual disability [39]. Male mice with Ptchd1 deletion exhibit attention deficits, gross hyperactivity, and poor distractor suppression during a visual detection task [40]. In mice, Ptchd1 expression is limited to the thalamic reticular nucleus during development, suggesting that an involuntary behavioral assay that indexed neural activity at higher stages of the central auditory pathway might be able to capture an auditory overload phenotype without the contribution of attentional and gross motor deficits that might otherwise complicate the interpretation of operant behavioral data.

ABR thresholds were equivalent in Ptch1 KO mice and wildtype controls, demonstrating that any differences in auditory responses were unlikely attributable to differences in early auditory processing. Quantitative videography revealed that the baseline pupil diameter was abnormally large in Ptchd1-KO mice, consistent with descriptions in several other strains of mice with autism risk gene mutations (**Figure 7B-C)** [41]. However, sound-evoked pupil dilation magnitude across a range of broadband noise levels was not different between KO and WT mice, after baseline pupil differences were factored out (two-way repeated measures ANOVA, N = 11/14; main effect for sound intensity [F = 17.74, p < 0.001]; main effect for genotype [F = 3.4, p = 0.08]; intensity × genotype [F = 2.04, p = 0.06]). By contrast, baseline FME was not significantly different between WT and KO mice (two-sample t-test, N = 11/15, p = 0.25; **Figure 7D**), but sound-evoked FME grew at a faster rate across a range of moderate sound intensities in KO mice compared to WT controls (**Figure 7E**), consistent with an auditory hyper-responsivity phenotype observed in persons on the autism spectrum as well as mice with acquired hyper-responsivity following sensorineural hearing loss (**Figure 5G**).

**Figure 7.**
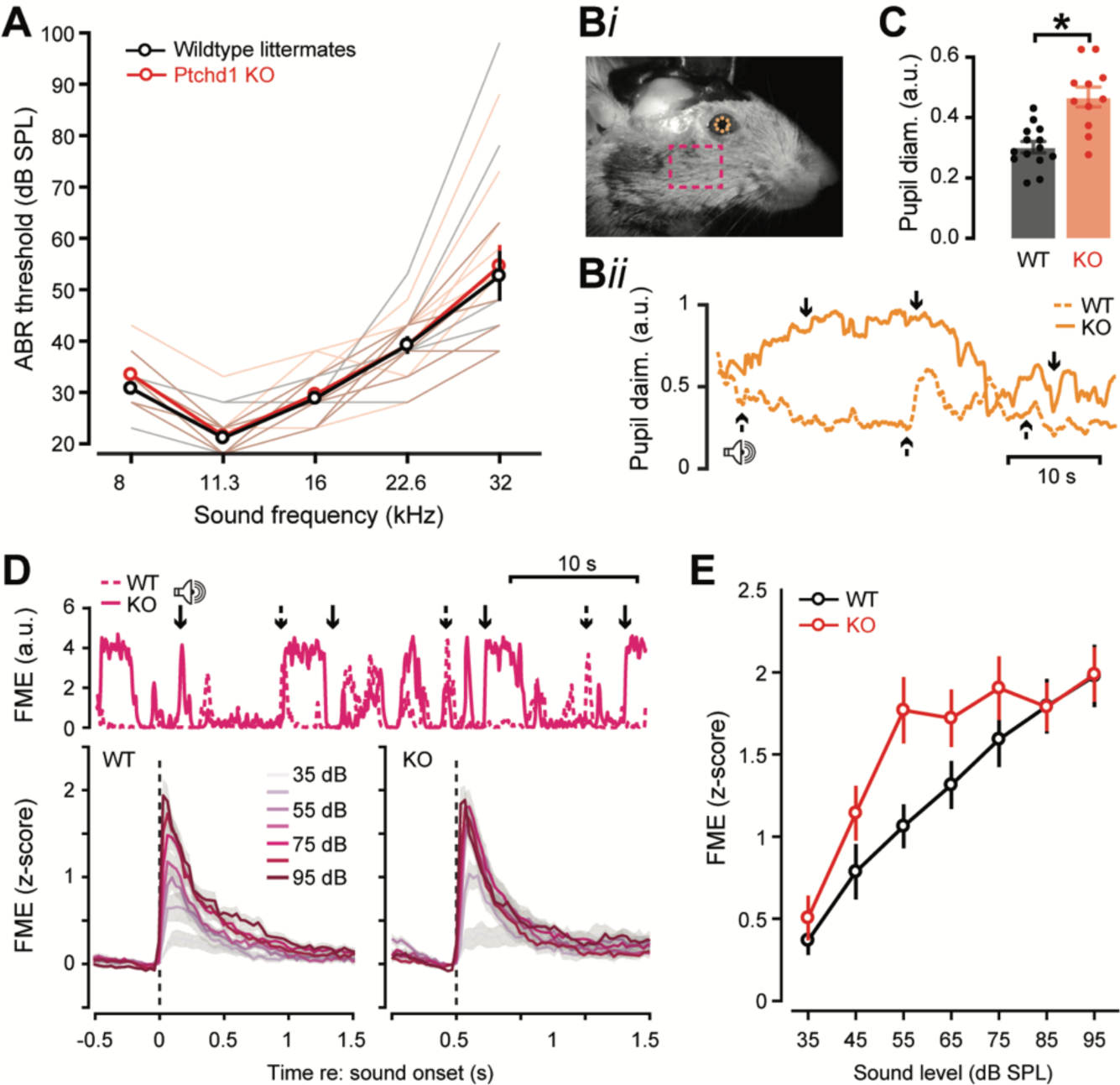
Hyper-responsive sound-evoked facial movements in mice with an autism risk gene mutation. (A) ABR thresholds are not significantly different in mice with Ptchd1 deletion (KO) and wildtype littermate controls (two-way repeated measures ANOVA, main effect for frequency [F = 131, p < 0.001], main effect for genotype [F = 0.31, p = 0.58], frequency × genotype interaction [F = 0.74, p = 0.60]). (B) *Bi:* Placement of pupil markers and ROI for FME calculation. *Bii*: Representative traces of pupil diameter changes in a wildtype (WT) and KO mouse. Solid and dashed arrows denote the timing of noise bursts in the corresponding recording session. (C) Baseline pupil diameter in KO mice is significantly larger than WT controls (two-sample t-test, [t = −4.64, p < 0.001]). Bars and errors bars represent mean ± SEM. Each data point represents an individual mouse (N = 14/11, WT/KO). (D) *Top:* Representative FME traces in a wildtype (WT) and KO mouse. Solid and dashed arrows denote the timing of noise bursts in the corresponding recording session. *Bottom:* Mean ± SEM FME for noise bursts of increasing intensity in WT (left) and KO (right) groups (N = 15/11, WT/KO). (E) Sound-evoked facial movement grow significantly more steeply across sound level in KO mice compared to WT controls (two-way mixed design ANOVA, main effect for sound level [F = 43.46, p = 0.001], main effect for genotype [F = 2.29, p = 0.14], level × genotype interaction [F = 2.67, p = 0.02]).

## DISCUSSION

We used quantitative videography to characterize sound-evoked movements of the face. We found that sound elicited small movements of a region of the face just caudal to the vibrissae array that was 30-40 dB more sensitive than the acoustic startle reflex and less variable than other points of measurement on the face (**Figure 1)**. FME faithfully encoded the low-frequency envelope of broadband stimuli up to 3 Hz (**Figure 2**), supporting accurate single-trial decoding of speech token identity in varying levels of background noise (**Figure 3**). We found that low-threshold sound-evoked FME was only elicited by sounds with spectral bandwidth greater than one octave (**Figure 4**) and showed no signs of habituation across repeated measurements over multiple weeks (**Figure 5)**. After noise-induced damage to the high-frequency cochlear base, FME was attenuated for a high-frequency stimulus, but low-frequency sounds targeting the spared region of the cochlea elicited hyper-responsive facial movements that were strongly correlated with potentiation of ABR wave 4, generated by the auditory midbrain. Firing rates of some deep layer A1 RS units were entrained by spontaneous facial movements, yet optogenetic A1 RS unit silencing potentiated sound-evoked FME, indicating that the auditory cortex is a modulator - not a mediator - of sound-evoked facial movements (**Figure 6**). Finally, we report that FME is hypersensitive at intermediate sound intensities in the Ptchd1-KO mouse, consistent with an auditory hypersensitivity phenotype associated with autism spectrum disorder (**Figure 7**).

### Facial videography is a sensitive readout of behavioral registration

Hearing, the conscious awareness of sound, is a psychological construct that can only be directly assessed through behavioral measurements. Operant assays for behavioral reporting of conscious sound awareness represent the gold standard for hearing assessments. However, training animals to perform operant tasks can take weeks, which makes measurements of complex sound processing difficult in most animal models and prohibitive for models of disordered sound perception that also have generalized motor, vestibular or cognitive impairments [45–48]. Alternatively, physiological proxies for hearing, such as the auditory brainstem response (ABR) can be measured in minutes but, as gross electrical potentials generated in the brainstem of anesthetized animals, these measures are not interchangeable with hearing assessments. For example, selective lesions of the cochlear afferent pathway render the ABR absent or grossly abnormal, yet auditory detection thresholds measured with operant behaviors are unaffected [19,23,49].

While sound-evoked facial movements provide high-fidelity readout of how sounds are registered by the central nervous system, the approach also has limitations. First, involuntary behaviors do not reflect conscious sound awareness, which can only be demonstrated with operant psychophysical behaviors. In addition, auditory research relies heavily on narrowband sounds, yet sound-evoked facial movements were largely insensitive to these stimuli. Despite this broadband sound preference, we demonstrated that frequency specific measurements using band-limited noise were possible and could report bidirectional changes following cochlear damage similar to operant behavior [21]. Sound-evoked facial movements also more easily disrupted by background noise than operant behavior, as FME-based stimulus decoding was significantly degraded at 20 dB SNR, while sound processing in noise studied with operant methods is generally robust down to SNRs well below zero [16,42,44,46]. Most importantly, perceptual awareness in operant tasks is closely linked to cortical registration of the sensory stimulus [16,48,50,52]. While ACtx is required for many forms of operant detection and discrimination [63–66], it is not required for sound-evoked FME. These observations underscore that while FME is relatively easy to measure and offers advantages over other behavioral and electrophysiological hearing indices, it is not necessarily a measure of perceptual sound awareness.

### The neural basis of sound-evoked facial movements

We found that optogenetic inactivation of ACtx paradoxically increased sound-evoked facial movements. This result suggests that the ACtx may modulate obligatory subcortical pathways for sound-evoked facial movements via the extensive network of descending corticofugal projections [36,43,45,47,49]. Given the strong correspondence between plasticity in the later waves of the ABR which are generated by the auditory midbrain [27,28] and FME at spared frequencies, our data suggest that sound-evoked facial movements are routed through the inferior colliculus, though the definitive evidence for this claim would require a series of lesions or inactivation experiments in various subcortical centers for sound processing. The possible involvement of the auditory midbrain is consistent with reports that electrical or optogenetic stimulation of the of the inferior colliculus induces immediate motor responses [45,51], including vibrissae movement in anesthetized rats [53], while optogenetic activation of the auditory thalamus is not known to induce motor activity [18,54]. Inferior colliculus neurons can be suppressed by electrically or optogenetically activating auditory cortex [55,56] through recurrent inhibition in intracollicular circuits [57], though the net effect of cortical stimulation varies with activation parameters [55,58]. Conversely, silencing ACtx increases spontaneous and sound-evoked activity in the inferior colliculus [59]. An increase in sound-evoked FME responses during cortical silencing is therefore consistent with the release of tonic inhibition imposed by corticollicular inputs.

While the afferent signals which result in sound-evoked facial movements may be routed through the auditory midbrain, it is clear from their sensory selectivity for broadband sounds that additional stages of sensory processing must occur in downstream areas, as virtually all nuclei within the central auditory pathway are responsive to pure tones at low sound intensities [60–62]. Similar selectivity for broadband sounds, but not pure tones, has been found in a reticular-limbic auditory pathway, though auditory response thresholds from neurons in these brain regions are higher than FME thresholds reported here [63].

Inferior colliculus neurons project primarily to the auditory thalamus, but also to the dorsal medial periaqueductal gray and the intermediate and deep layers of the superior colliculus [45]. The dorsal periaqueductal gray is known to be involved in the generation of defensive behaviors [64,65], but defensive behaviors elicited by sound should rapidly adapt within one to two stimulus presentations [66], inconsistent with the indefatigable sound-evoked facial movements we observed. In contrast, the intermediate and ventral layers of the superior colliculus are known to respond robustly to sound, with receptive fields that are preferential towards high frequency sounds with large spectral bandwidth [67,68]. These high-frequency, large spectral bandwidth receptive fields are congruent with our finding that baseline FME responses were greater for an octave band noise centered at 32 kHz compared to 8 kHz. Further, anatomical tracing has shown that the superior colliculus projects directly to the motoneurons of the facial nucleus of the mouse [69], and optogenetic activation of the SC elicits attempted head-movements in head-fixed mice [70]. It remains possible that sound-evoked facial movements could be routed through a brainstem pathway, as it is clear that audiomotor pathways have not been exhaustively characterized. For instance, a novel multi-synaptic pathway from the lateral lemniscus in the auditory brainstem to anterior lateral motor cortex has recently been described [71]. Further work will be required to elucidate the precise pathway through which auditory signals evoke facial movements.

### Broader consequences of sound-evoked facial movements for neuroscience research

A broad class of auditory inputs elicit a short-latency cascade of motor efferent and somatosensory reafferent activity in the brain. The implication of this finding is that within approximately 30 ms after sound presentation, activity in many brain regions of awake mice will reflect a mixture of auditory, motor, and somatosensory activity. Disentangling the relative weighting of auditory, motor, or somatosensory contributions to multisensory interactions, where activity between sensory systems could intermix at various levels of sensory processing and timescales, has proved challenging. For example, a recent study demonstrated that auditory influences on visual cortex response properties could mostly be accounted for by global, low-dimensional activity patterns arising from sound-evoked facial movements [12]. “Unisensory” experiments must also reckon with the influence of motor corollary and sensory reafferent inputs. We reported here the spiking rates of a substantial minority of regular spiking units in layers 5 and 6 of A1 were significantly modulated shortly after spontaneous facial movements, supporting earlier mesoscale recordings that described widespread activity across the dorsal cortex related to facial movements [72]. We [73] and others [34] have shown that planned movements of the face or forelimb modulate the spiking of layer 5 and 6 neurons hundreds of milliseconds prior to movement and sound onset, highlighting the inextricable link between movement and sound and underscoring the difficulty of studying one in the absence of the other, at least in awake subjects.

However, our work shows that facial movements are not elicited by all sounds, but rather are only elicited by broadband stimuli. Thus, by carefully controlling the stimulus (i.e. using narrowband sounds presented at moderate sound pressure levels), potential confounds caused by sound-evoked facial movements may be mitigated if not avoided altogether.

### Sound-evoked facial movements as a tool for active sensing

Most reflex pathways have a clear adaptive value to the animal, though the adaptive benefit of sound-evoked facial movements is not entirely clear. For example, the acoustic startle reflex has been suggested to not only to protect the body from immediate harm [74], but also to rapidly terminate ongoing sequences of motor activity and prepare escape behaviors [75]. One possibility is that sound-evoked facial movements may be a component of orienting behaviors [70,76]. Complementary to moving the head towards a sound source, moving the face could allow the mouse to actively sample its environment using its vibrissae arrays. The sensory tuning of facial movements we describe both matches the fact that most natural sounds are broadband [77] and that broadband sounds provide robust interaural level difference cues for sound source localization and phonotaxis [68]. Future work using head-mounted cameras in freely-moving animals [11,78,79] could clarify the relationship between orienting responses and sound-evoked facial movements.

## Acknowledgments

We thank Dr. Michael Halassa for the gift of the Ptchd1-KO mouse. We thank Ashvini Melkote for assistance with DeepLabCut, Dr. Meenakshi Asokan for providing videography analysis code and performing initial pilot experiments, Dr. Matthew McGill for assistance with cochlear function measurements, and Divya Narayanan and Yurika Watanabe for providing surgical support.

## Author contributions

Conceptualization, K.K.C., K.C. and D.B.P.; Methodology, K.K.C, A.G., K.C., K.E.H.; Investigation, K.K.C, K.S.S., A.R.C., and K.C.; Software, K.K.C., A.G., A.R.C, K.C and K.E.H.; Formal Analysis, K.K.C., A.R.C, K.C., and A.G.; Data Curation, K.K.C., A.R.C, K.C., A.G., and K.S..; Visualization, K.K.C., A.R.C, K.C. and D.B.P., Writing – Original Draft, D.B.P. and K.K.C.; Writing – Review & Editing, K.K.C. and D.B.P; Resources, D.B.P; Supervision, D.B.P.; Funding Acquisition, D.B.P.

## STAR ✭ Methods

### RESOURCE AVAILABILITY

#### Lead Contact

Further information and requests should be directed to and will be fulfilled by the lead contact, Kameron Clayton.

#### Materials Availability

This study did not generate new reagents.

#### Data and Code Availability

Data acquisition and analysis was performed with custom scripts in MATLAB, LabVIEW, and Python. Spike sorting was done in Kilosort 2.0 (https://github.com/cortex-lab/Kilosort). Markerless behavior tracking was done with DeepLabCut (https://github.com/AlexEMG/DeepLabCut). Data associated with this article was deposited in Mendeley Data (DOI: TBD). Original code is publicly available (github.com/TBD) and further information required for re-analysis of data reported in this paper is available from the lead contact upon request.

#### Animal subjects

We used 76 adult male and female mice aged 8-12 weeks. Mice were maintained on a 12/12 hr light/dark cycle and all experiments performed during the dark cycle, with food and water available ad libitum in the home cage. Mice were housed individually following major survival surgery. All procedures were approved by the Massachusetts Eye and Ear Animal Care and Use Committee and followed the guidelines established by the National Institute of Health for the care and use of laboratory animals.

For simultaneous videography and acoustic startle testing, 8 C57 × Cdh23 mice were used. C57 mice were crossed with homozygous Cdh23 mice to prevent the precocious high frequency hearing loss typically found in C57 mice, which can begin as early as 12 weeks post-natal [80]. For videography experiments with temporally modulated stimuli, an additional 8 C57BL6 × Cdh23 mice were used. For noise exposure experiments, we used a total of 20 C57BL6 × Cdh23 (10 exposed/8 sham). Auditory brainstem responses were collected in a subset of 9 noise-exposed mice, with one mouse excluded from further analysis because of the absence of wave IV in pre-exposure measurements. Single-unit electrophysiology was performed in 5 C57BL6 × Cdh23 mice. Videography with optogenetic stimulation was performed in 7 PV-Cre × Ai32 mice and 3 PV-Cre mice with injected with virus to express channelrhodopsin bilaterally in ACtx. Experiments characterizing sound-evoked facial movements in the Ptchd1-KO mouse were performed in 11 KO mice and 15 C57 littermate controls.

### METHOD DETAILS

#### Surgical preparation

Mice were anesthetized with isoflurane in oxygen (5% induction, 1.5-2% maintenance). Body temperature was maintained at 36.6 with a homeothermic blanket system (FHC). Lubricating ointment was placed on the eyes. Lidocaine hydrochloride (0.1 mL) was administered subcutaneously to numb the scalp. For analgesia, Buprenex (0.05 mg/kg) and meloxicam (0.1 mg/kg) were administered subcutaneously at the beginning of the procedure and again 24 and 48 hours from the initial dosing. Following surgery, the mice were transferred to a heated recovery chamber.

Following repeated serial applications of Betadine and 70% ethanol, the skin overlying the dorsal cranium was retracted and the periosteum was removed. Etchant (C&B metabond) and 70% ethanol was applied to prepare the exposed skull surface. For mice not undergoing optogenetics experiments, a custom titanium headplate (iMaterialise) was affixed to the skull with dental cement (C&B metabond).

#### Optical access to auditory cortex

To prepare mice for bilateral optogenetic stimulation, the skull overlying each auditory cortex was made optically transparent. First, a layer of super glue (Krazy-Glue) was applied and allowed to dry, followed by layers of clear cement (C&B metabond) and nail polish (L.A. colors). While the nail polish dried, black plastic casings (Freelin-Wade) were affixed around the transparent portion of the skull to allow for interfacing with optic fiber patch cables. Once the casings were affixed, a custom titanium headplate was affixed to the dorsal surface of the skull with cement.

For PV-Cre mice only, channelrhodopsin was expressed in PV neurons by injecting AAV5-EF1a-DIO-ChR2 into ACtx. Briefly, two small burr holes were made in the skull overlying ACtx in each hemisphere using a 31-gauge needle, 1.5-2.25 mm rostral to the lambdoid suture. The viral solution was backfilled into a pulled glass pipette (Drummond, Wiretrol II) and a precision injection system (Drummond, Nanoject III) was used to deliver 200 nL of virus per injection site at a rate of 9 nL/min, 0.3 mm below the pial surface. The pipette was left to rest for at least 10 minutes following the end of the injection and burr holes were filled with KWIK-SIL (WPI). Injections took place immediately prior to the skull clearing procedure.

#### Videography

A CMOS camera (Teledyne Dalsa, model M2020) equipped with a lens (Tamron 032938) and a long-pass filter (MidWest Optical, LP830, 25.5nm cutoff) was positioned approximately 25 cm to the right of the animal’s face, and illuminated by an array of infrared LEDs (Vishay, 850 nm). A white LED (Thorlabs, MBB1F1) was used to provide ambient illumination sufficient to keep the pupil diameter in an intermediate range. Video recordings of the face and foot were acquired with a 512 × 512 pixel resolution at 150 Hz (for data shown in Figures 1-5) or 30 Hz (for data shown in Figures 6-7).

Auditory stimuli were generated with a 24-bit digital-to-analog converter (National Instruments model PXI-4461), amplified (Samson,120a Power Amplifier), and presented via a tweeter (ScanSpeak) positioned 25 cm from the left ear. Speaker output calibrated with a 1/4” prepolarized microphone (PCB Electronics).. Mice were head-fixed atop an acrylic plate resting on three piezoelectric force transducers (PCB Piezotronics) coupled to a summation amplifier, allowing measurement of the downward force caused by skeletal muscle contraction. Mice were allowed to acclimate for 15 minutes before recordings began.

#### Stimulus generation and presentation

Each stimulus was repeated in 20 trials in a pseudo-random order separated by an intertrial interval duration selected at random from a truncated exponential distribution to produce a flat hazard function. Broadband noise bursts (50 ms duration, 5 ms raised cosine onset and offset ramps) were presented between 15 to 115 dB SPL in 10 dB steps with a 10-20s intertrial interval. For gaps in noise, silent gaps (30 - 500 ms in duration, 0.1ms onset and offset ramps) were inserted in 50 dB SPL continuous broadband noise with a 14 - 19 s inter-trial interval. For FM sweeps, sequences of 6 sweeps (200 ms duration, ±20 octaves/second between 4-64 kHz, 5 ms raised cosine ramps) were presented at 1 - 4 Hz in 0.5 Hz steps with a 7 - 11 s intertrial interval. Speech stimuli were 200 ms tokens produced by an adult female speaker resynthesized to be four octaves higher than the original source material with the TANDEM-STRAIGHT vocoder [81], as originally synthesized by Chambers et al. [20]. Speech tokens (70 dB SPL) were presented in trains of 6 tokens at 1 Hz in continuous broadband noise (10 - 50 dB SPL in 10 dB increments) or in silence, with a 7-11s intertrial interval. To contrast tone and broadband noise responses, tone bursts (50 ms duration with 5 ms raised cosine ramps at 8 and 32 kHz or broadband noise bursts were presented from 30 - 90 dB SPL in 20 dB increments with a 7 to 11 s intertrial interval. Octave-band noise bursts were made by applying a 4^th^ order Butterworth filter centered at 8 or 32 kHz to broadband noise. Octave-band noise bursts (50 ms duration with 5 ms raised cosine ramps) were presented from 30 - 100 dB SPL in 10 dB increments with a 7 - 11 s inter-trial interval.

#### High-frequency noise exposure

To induce high-frequency SNHL, octave-band noise at 16 - 32 kHz was presented at 103 dB SPL for 2 hours. Exposure stimulus was delivered via a tweeter fixated inside a custom-made exposure chamber (51 × 51 × 51 cm). The interior walls of the acoustic enclosure joined at irregular, non-right angles to minimize standing waves. Additionally, to further diffuse the high-frequency sound field, irregular surface depths were achieved on three of the interior walls by attaching stackable ABS plastic blocks (LEGO). Prior to exposure, mice were placed, unrestrained, in an independent wire-mesh chamber (15 × 15 × 10 cm). This chamber was placed at the center of a continuously rotating plate, ensuring mice were exposed to a relatively uniform sound field. Sham-exposed mice underwent the same procedure except that the exposure noise was presented at an innocuous level (70 dB SPL). All sham and noise exposures were performed at the same time of day.

#### Cochlear function testing

Animals were anesthetized with ketamine (120 mg/kg) and xylazine (12 mg/kg), placed on a homeothermic heating blanket during testing, and administered half the initial ketamine dose as a booster when required. Acoustic stimuli were presented via in-ear acoustic assemblies consisting of two miniature dynamic earphones (CUI CDMG15008–03A) and an electret condenser microphone (Knowles FG-23339-PO7) coupled to a probe tube. Stimuli were calibrated in the ear canal in each mouse before recording. The ABR was measured with tone bursts (5 ms with 0.5 ms raised cosine ramps at 8,11.3,16, 22.6, and 32 kHz delivered at 26.99 Hz from 20-100 dB SPL in 5 dB increments) and the same octave-band noise bursts described above delivered at 10.02 Hz. Recordings were made with one transdermal electrode behind the right pinna and one electrode attached to a silver wire (A-M Systems) placed on the surface of the brain at vertex through a burr hole and cemented in place during the initial headplate surgery. Threshold was determined as the lowest intensity which elicited a repeatable waveform. Positive and negative peaks of each ABR wave were quantified as the peak to trough amplitude of each wave, subtracted by the peak to trough amplitude of the pre-stimulus baseline to correct for the measurement noise floor. ABR testing was performed 1 week before noise or sham exposure and again two weeks after exposure, following the final behavioral measurement.

#### Cortical electrophysiology

##### Preparation for acute insertion of high-density probes in awake, head-fixed mice

A ground wire was placed over the left occipital cortex through a small burr hole during the initial head plate attachment surgery. On the day of recording, the mouse was briefly anesthetized with isoflurane in oxygen (5% induction, 2% maintenance) and a scalpel was used to make a small (1 × 1 mm) craniotomy over right auditory cortex, centered on the temporal ridge between 1.5 and 2.5 mm anterior from the lambdoid suture. A well was constructed around the craniotomy using UV-cured composite (Flow-It ALC) was filled with lubricating ointment (Paralube Vet Ointment), after which isoflurane was discontinued and the mouse was moved to a body cradle where the animal’s head was immobilized by attaching head plate to a fixation post in a dimly lit double walled acoustic chamber. Recordings began 30 minutes after the cessation of isoflurane to allow for full recovery from anesthesia. Following each experiment, the recording chamber was flushed with saline, lubricating ointment was reapplied, and capped with UV-cured composite.

##### Extracellular recordings

A single shank 64- channel silicon probe (Cambridge Neurotech: H3, 20 μm between contacts) was positioned perpendicular to the cortical surface using a micromanipulator (Narishige) and inserted using a hydraulic micromanipulator (Narishige). The probe was advanced at 100 μm/s until the probe tip was approximately 1.3-1.4 mm below the cortical surface. The probe was allowed to settle for 15 minutes before recordings began. At the beginning of each recording, noise bursts (50 ms duration, 70 dB SPL, 50 repetitions) were presented to confirm the depth of the probe relative to auditory cortex using the current source density [33,82]. Additionally, frequency-receptive fields were derived to confirm the placement of the probe within the primary auditory cortex, using the tonotopic reversal which marks the rostral border of A1 [61]. We simultaneously recorded facial videography and laminar electrophysiology as mice were presented with frequency-modulated sweeps (4-64 kHz, 80 and −80 oct/s, 50 ms duration, 30-70 dB SPL in 10 dB increments, 60-80 repetitions) from a free-field speaker placed approximately 10 cm from the left ear.

#### Optogenetic stimulation

Neurons transduced with channelrhodopsin were activated through the optically cleared skull with light (10 mW at fiber tip, 500 ms, 20 Hz pulse rate, 25 ms pulse width, terminating with 100 ms linearly decreasing ramp) that have been shown to trigger robust inhibition of pyramidal neuron spiking without rebound excitation [83]. Blue light was delivered by two 473 nm diode lasers (Omicron LuxX) via optic fibers terminating in ferrules (Doric, 0.2 mm diameter, 0.22 NA), which were fit with mating sleeves to create a snug interface with the skull-mounted plastic casings. Laser onset preceded sound onset by 100 ms. Laser stimulation occurred in half of trials, which consisted of 50 ms noise bursts presented from 35 to 95 dB SPL in 10 dB increments, with a 6-10 s inter-trial interval.

### QUANTIFICATIONS AND STATISTICAL ANALYSIS

#### Video processing and analysis

##### Facial movement energy

A region of interest (ROI) was manually drawn on the rostral cheek, just caudal to the vibrissae array. FME was defined as the sum of the absolute difference in intensity between consecutive frames for each pixel within the ROI [84,85]. FME was z-scored with respect to the mean and standard deviation of the session.

##### Markerless behavior tracking

DeepLabCut was used to track the nose, ear, jaw, hindpaw, eyelid diameter, and pupil diameter. The anterior tip of the nose, posterior edge of the pinna, jaw, and the metatarsal joint of the hindfoot were labeled with single markers. The eyelid and the pupil were labeled with 8 markers that spanned the four cardinal and four intracardinal compass points. Separate DeepLabCut models were used for points on the face and the foot. For the face model, 300 frames from 30 mice were used to train the model (model originally used in Robert et al. [86]). For the foot model, 40 frames from 6 mice were used. For both models, a ResNet-101 based neural network was trained on 95% of the labeled frames for 1,030,000 training iterations, using the default parameters in DeepLabCut.

Any time point for which model tracking likelihood dropped below 90% was discarded and interpolation was performed between neighboring frames. For single-point tracking, movement amplitudes were taken as the square root of the sum of the squared × and y velocities. For eyelid and pupil diameter, tracked points were used to fit an ellipse using a least-squares criterion and calculating the long axis diameter. Because the eyelid was widest at rest and narrowed with sound stimulation, we took the absolute value of the eyelid diameter so that movement would be positive going. All movement traces were z-scored with respect to their mean and standard deviations for the session.

##### Quantification of sound-evoked movements

Sound-evoked response amplitude for all movements were taken as the mean of the 5 frames surrounding the peak response within 1 second of stimulus onset. To determine the threshold of sound-evoked movement, we performed a paired t-test comparing the mean of 1 s pre-stimulus and the post-stimulus response across trials. Threshold was defined as the lowest intensity at which reliably elicited a significant response; all levels above threshold were required to have a significant response. Latency was defined as time to the half-maximum of the response. Intertrial variance was defined as the coefficient of variation, the standard deviation of trial-by-trial sound-evoked responses divided by the mean response for each subject individually. As computing the coefficient of variation for intersubject variance across subjects would give a single value for each measured effector, we treated each trial as an individual measurement and computed the standard deviation of the sound-evoked response divided by the mean response across subjects for each trial separately.

To quantify the rhythmic entrainment of facial movements, we used the fast Fourier transform to compute the power spectral density (PSD) within the stimulus period, which varied in duration depending on the repetition rate from 1.5 to 6 seconds. The dB SNR metric was computed as 10 times the common logarithm of the PSD amplitude at the stimulus repetition rate divided by the average PSD amplitude at neighboring frequencies. To decode syllable identity trial by trial, a support vector machine with a linear kernel was trained for each mouse using time series responses to each syllable in quiet. All data was first transformed using principal components analysis and only the principal components which cumulatively explained 80% of the variance were included to prevent overfitting. We used 10-fold cross-validation to train and test the classifier and repeated this process for 1000 iterations using different random train/test splits. To estimate the classification noise floor, we shuffled the trial-by-trial syllable labels and repeated the training and testing process. Models were fit using the Matlab function ‘fitcsvm’. For FME response tracking over time, fold change was calculated as the response for each day divided by the baseline response, which included the two pre-exposure days, for 0 and 10 dB re: threshold.

#### Acoustic startle reflex measurement

The acoustic startle reflex is highly-stereotyped triphasic signal which corresponds to the contraction of skeletal muscles [13]. We quantified the startle reflex amplitude as the peak to trough of the response in the 1 s following stimulus onset and computed measures of threshold, latency and variance as we did for facial videography.

#### Electrophysiology acquisition and online analysis

Raw neural signals were acquired at 24.4 kHz and digitized at 32-bit resolution (PZ5 Neurodigitizer, RZ2 BioAmp Processor, RS4 Data Streamer; Tucker-Davis Technologies). For online data visualization, the common mode, channel-averaged, signal was subtracted from the raw signals for all channels to eliminate artifacts shared across all channels. To examine multiunit activity, signals were band-pass filtered (300-3000 Hz, second-order Butterworth filters) and the threshold for spike detection was set as a negative deflection greater than 3.5 standard deviations above the background hash. Following notch filtering at 60 Hz and downsampling to1 kHz, the CSD was computed as the second spatial derivative of the local field potential.

Signals were spatially smoothed across channels using a 5-point Hanning window to mitigate potential artifacts induced by impedance mismatches between neighboring channels. The layer 4/5 boundary (0.5 mm from the pial surface) was identified using the noise-evoked CSD [33,82].

#### Single unit identification and analysis

Kilosort 2.0 was used to sort spikes into single unit clusters [87]. Single-unit classification was based on the presence of a well-defined refractory period in the interspike interval histogram and an isolation distance (>10) which indicated that the cluster was well separated from noise [88]. Units were classified as RS if the peak-to-trough delay of its mean spike waveform was greater than 0.6 ms.

#### Statistical analysis

All statistical analysis was performed in Matlab. We used the standard p < 0.05 threshold for assigning statistical significance. Multiple post-hoc comparisons were corrected with the Holm-Bonferroni correction.

